# Capturing brain dynamics: latent spatiotemporal patterns predict stimuli and individual differences

**DOI:** 10.1101/2020.06.11.146969

**Authors:** Manasij Venkatesh, Joseph JaJa, Luiz Pessoa

## Abstract

Insights from functional Magnetic Resonance Imaging (fMRI), and more recently from recordings of large numbers of neurons through calcium imaging, reveal that many cognitive, emotional, and motor functions depend on the multivariate interactions of neuronal populations. To capture and characterize spatiotemporal properties of brain events, we propose an architecture based on long short-term memory (LSTM) networks to uncover distributed spatiotemporal signatures during dynamic experimental conditions^1^. We demonstrate the potential of the approach using naturalistic movie-watching fMRI data. We show that movie clips result in complex but distinct spatiotemporal patterns in brain data that can be classified using LSTMs (≈ 90% for 15-way classification), demonstrating that learned representations generalized to unseen participants. LSTMs were also superior to existing methods in predicting behavior and personality traits of individuals. We propose a dimensionality reduction approach that uncovers low-dimensional trajectories and captures essential informational properties of brain dynamics. Finally, we employed saliency maps to characterize spatiotemporally-varying brain-region importance. The spatiotemporal saliency maps revealed dynamic but consistent changes in fMRI activation data. We believe our approach provides a powerful framework for visualizing, analyzing, and discovering dynamic spatially distributed brain representations during naturalistic conditions.

## 1 Introduction

As brain data become increasingly spatiotemporal, there is a great need to develop methods that can effectively capture how information across space and time combine to form representations of mental events supporting behavior. Although fMRI data are acquired temporally, they are most often treated in a quasi-static manner. However, a fuller understanding of the mechanisms that support mental functions necessitates the characterization of dynamic properties. To address this gap we describe methods to characterize both high- and low-dimensional *trajectories* that provide “signatures” for experimental conditions (Fig. 1B). We investigated them at the between-participant level (in contrast to within-participant) to ascertain the generalizability of the representations created by the approach. Although our methods can be applied to any kind of fMRI data, we focused here on data acquired during movie-watching given its inherent dynamics. We address the following questions: (1) Can brain signals generated by dynamic stimuli be characterized in terms of generalizable spatiotemporal patterns? How are such patterns distributed across space and time? (2) Understanding the dimensionality of brain representations has become an important research question in recent years ([1, 2, 3]). Accordingly, we sought to investigate the prediction accuracy of both high- and low-dimensional (3) If spatiotemporal patterns capture important properties of brain dynamics, do they capture information about individual differences that are predictive of an individual’s behavioral capabilities and/or personality?

**Figure 1:**
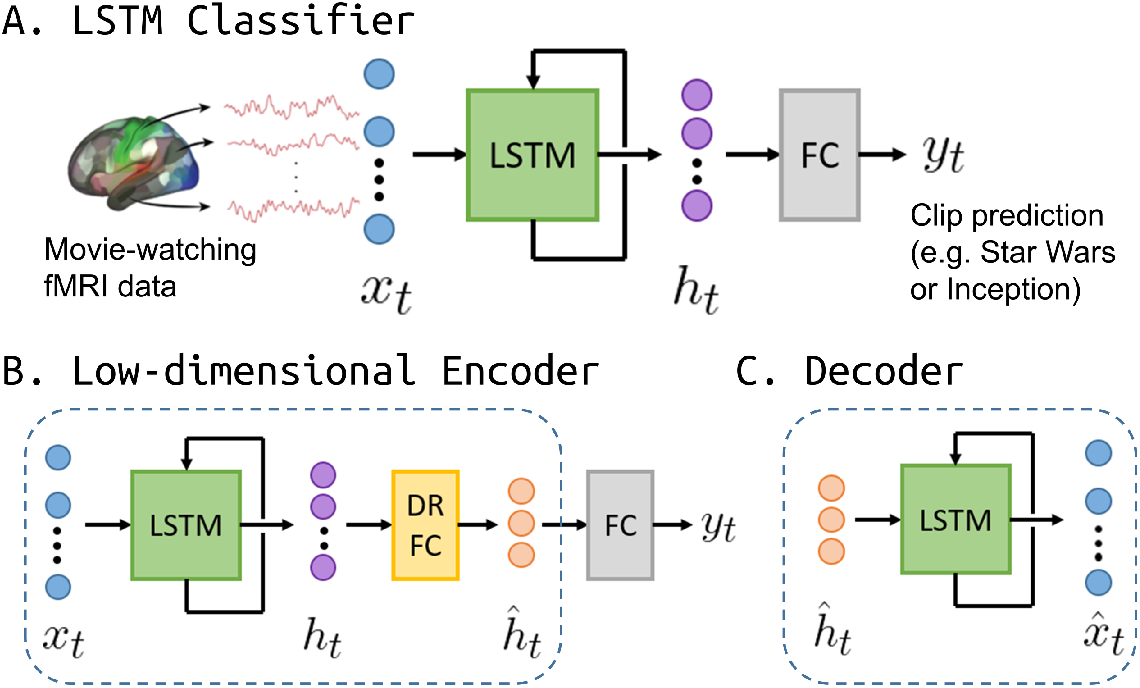
(A) The classifier consisted of Long Short-Term Memory (LSTM) units with a fully-connected (FC) dense layer for label prediction at each time step. (B) For dimensionality reduction, the LSTM outputs were first linearly projected to a lower dimensional space using a fully-connected layer (DR-FC). Classification was then performed on the low-dimensional representations, 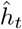. (C) LSTM decoder used to reconstruct the original time series from the low-dimensional representation.

## 2 Methods

We employed Human Connectome Project (HCP; [4]) movie-watching data. Participants were scanned while they watched movie excerpts (Hollywood and independent films) and other short clips (see Figure 3 legend for details), which we call “clips”. Data were sampled every 1 second. All 15 clips contained some degree of social and affective content. Participants viewed clips once, except for the *test-retest clip* that was viewed 4 times. We used all movie-watching HCP data, except for 8 participants with runs missing; thus we used *N* = 176. The preprocessed HCP data included FIX-denoising, motion correction, and surface registration (details in [5, 6]). We analyzed data at the region of interest (ROI) level, with one time series per ROI (average time series across spatial locations within ROI). We employed a 300-ROI cortical parcellation [7]. ROIs were also grouped based on large-scale network definitions ([8]; see Fig. 4).

### 2.1 Long Short-Term Memory for classification of brain data

Deep neural networks (DNNs) can be used for limited temporal modeling by means of sliding windows. Recurrent NN (RNN) architectures contain feedback cycles that allow signals at the current time step to be influenced by long-term information. However, training traditional RNNs is challenging give, for example, the problem of vanishing/exploding gradients that prevent them from learning relationships beyond 5-10 time steps [9]. The gating mechanisms in Long Short-Term Memory (LSTM) networks overcome these problems [10]. Here, we employ an LSTM-based architecture to characterize spatiotemporal structure in fMRI movie data.

Brain activation at time step, *x*_*t*_, was fed sequentially to an LSTM (Fig. 1A). The cell state *c*_*t*_ and hidden state *h*_*t*_ were updated as follows:

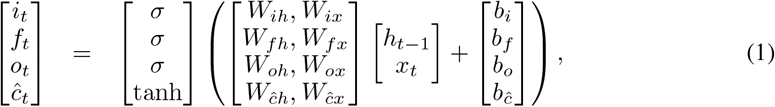

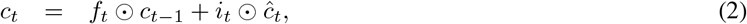

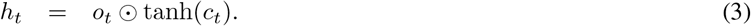

where the input gate, *i*, controls the extent to which the current input is exposed to the cell state; the forget gate, *f*, discards prior history; and the output gate, *o*, controls how the cell state affects gate activations during the next time step [11]. We represent the input size (number of ROIs) by *N*_*x*_ and the hidden state size by *N*_*h*_; other gates also have size *N*_*h*_. The *W* matrices contain the weights. For example, *W*_*ix*_ is *N*_*h*_ × *N*_*x*_ and represents the connections between the inputs, *x*, and the input gate, *i*. The bias weights are indicated by *b*, and *σ* represents the sigmoid activation function. The operator ⊙ represents element-wise multiplication. Each output of the LSTM, *h*_*t*_, was input to a fully-connected (FC) layer used to predict the input label (here, movie clip), *y*_*t*_ as

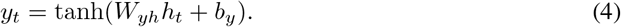

The dimensionality of *y*_*t*_ was 15 to implement 15-way classification based on the number of clips. Note that the output of the network, based on max *y*_*t*_, corresponds to a time series of label predictions. We analyzed classification accuracy as a function of time to identify when clips became separable in the latent space of hidden states. We refer to the LSTM together with the FC layer as the *LSTM classifier*. The classifier was trained using the Backpropagation Through Time algorithm minimizing the cross-entropy loss between the predictions and the true labels with PyTorch [12].

Data from 100 participants were used for training, and the remaining 76 participants were used for testing. To determine optimal hyperparameters for clip prediction, we employed a 10-fold cross-validation approach on the training data. In each fold, participants in the training set were not included in the validation set and vice versa.

### 2.2 LSTM-based dimensionality reduction

Unsupervised dimensionality reduction (DR) techniques such as Principal Component Analysis (PCA) project data onto a lower dimensional space such that variance is maximized. Since they do not account for class labels, representations may not be separable in the lower dimensional space. Supervised techniques such as Linear Discriminant Analysis ensure that in the projected space, data are most separable using a linear decision boundary. In most DR settings, samples are assumed to be statistically independent and thus do not account for the temporal relationships in time series data. Further, unless kernel methods are used, only linear mappings of the original high-dimensional data are possible. Here, we propose a non-linear supervised dimensionality reduction technique for temporal data.

LSTM outputs are typically high-dimensional (*N*_*h*_). Several researchers have proposed “probing” into intermediate layers for increased interpretability [13, 14]. Whereas these techniques have improved understanding of how representations are learned, here we probe into LSTM hidden states to visualize dynamics. LSTM outputs, *h*_*t*_, were linearly projected onto a lower dimensional space, 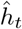, using an FC layer. We refer to this weight matrix of size 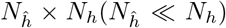 as the *Dimensionality Reduction Fully-Connected (DR-FC)* layer, and to this model as the *LSTM encoder*. Since 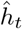 is a low-dimensional representation of the history of *x*_*t*_, the inputs are not treated independently, thus effectively leading to non-linear temporal dimensionality reduction. A final FC layer was used to predict labels based on 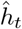.

### 2.3 LSTM decoder

Can low-dimensional representations obtained for classification be used to reconstruct the original data? We used an *LSTM decoder* to reconstruct *x*_*t*_ from 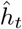 (Fig. 1C). The decoder was trained separately from the LSTM encoder, and minimized the mean squared error (MSE) loss between the LSTM decoder output 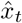 and the input *x*_*t*_. Note that the approach is different from autoencoders where latent representations are obtained such that the reconstruction loss is minimized. Here, the encoder training was independent from the decoder. To assess performance, we computed the fraction of the variance in the original data captured after reconstruction from low-dimensional representations and compared it to that captured by reconstruction from the same number of principal components.

### 2.4 Saliency maps for spatiotemporal importance

To understand the importance of each ROI at a given time step to clip prediction, we used saliency maps [15]. For each participant’s clip data, the gradient of the class score with respect to the input was computed by backpropagation. To determine ROIs that were consistently important across participants, we obtained mean saliency maps for each clip by averaging across test participants.

### 2.5 Baseline models

#### Feed-forward network

We also trained a deep feed-forward (*FF classifier*) network consisting of fully-connected layers. Each time step in the input time series was classified independently, and thus no temporal structure was modeled. The optimal number of layers was chosen based on cross-validation following a grid search between 2 and 10 layers. The number of units in each layer was the same as the hidden state size of the LSTM.

#### Temporal Convolutional Network

LSTMs model temporal dependency using a dynamically changing contextual window over the input time series. Are static fixed-sized windows sufficient for modeling dynamics in brain data? Thus, we employed a Temporal Convolutional Network (*TCN*; [16]), which maps an input time series of a given temporal length to an output time series of the same length (similar to LSTMs). However, convolutions are causal, such that the output at time *t* is based on convolutions only with inputs from time *t* and earlier. A fully-connected layer was used to predict clip labels based on TCN outputs, which we call *TCN Classifier.* To investigate different temporal windows, we varied the kernel width from 10 to 50 in steps of 10. By fixing the kernel height to the number of ROIs, we ensured that convolutions were only along the temporal dimension, and each kernel resulted in a 1D time series. For comparisons with LSTMs, the number of kernels was set to the optimal hidden state size of the LSTM, so that the output of the TCN was also of size *N*_*h*_.

### 2.6 Predicting behavior and personality traits

LSTMs were also trained to predict behavior and personality-related scores: fluid intelligence, verbal IQ, and personality measures [17]. Using clip data as input, an LSTM model was trained to predict behavioral scores at each time step by minimizing the MSE loss between the predictions *y*_*t*_ (a single unit with a continuous-valued output) and the true scores.

## 3 Results

### 3.1 Generalizability of spatiotemporal patterns in naturalistic fMRI data

To understand whether watching specific clips result in spatiotemporal brain patterns that are generalizable across participants, we employed the *LSTM classifier* to predict clip labels. The optimal hidden state size was set to *N*_*h*_ = 150 based on cross-validation (we did not find improvements using larger sizes). Classification accuracy was 87.35% for 15-way classification (Fig. 2A). Since the outputs of the classifier were clip predictions at each time step, we also analyzed true positive rate (TPR) as a function of time for each clip (Fig. 2B). For most clips, TPR was poor at the beginning of the clip but increased sharply for the first 30 seconds, and then gradually reached over 90%. A formal evaluation of chance performance based on the null distribution obtained through permutation testing (1000 iterations; [18]) resulted in a mean null accuracy of 6.67%.

**Figure 2:**
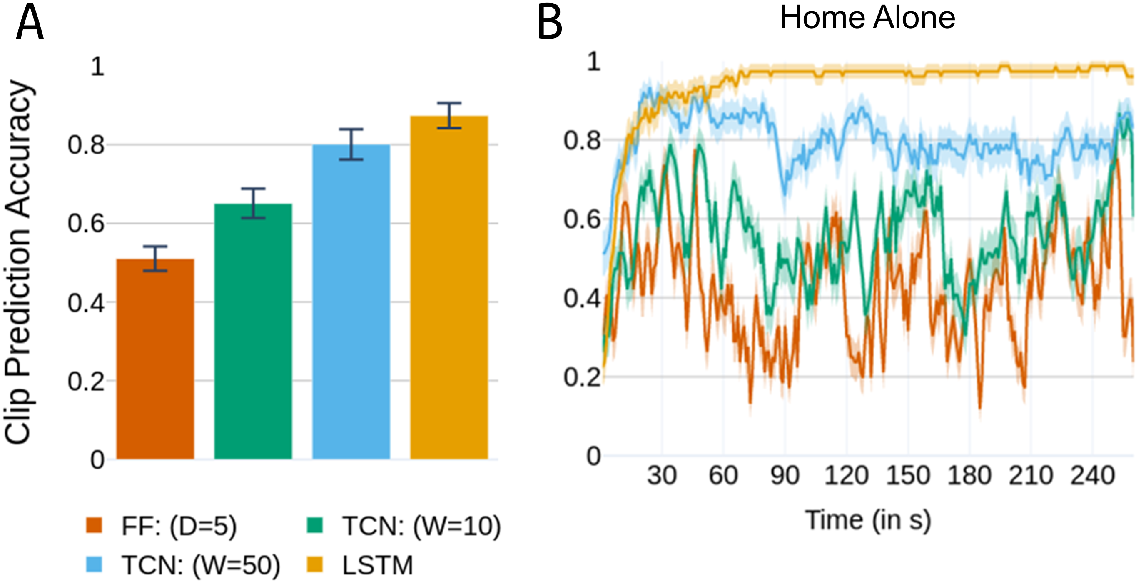
LSTMs and competing models. (A) Mean clip prediction accuracy (15-way classification) using feed-forward (FF, 5 layers) classifiers, temporal convolutional networks (TCN, kernel widths of 10 and 50), and LSTMs. (B) Since our framework predicted labels at each time step, true positive rate for each clip was determined as a function of time (illustrated for *Home Alone*). Error bars show the standard error of the mean across test participants.

### 3.2 Is temporal information necessary for clip prediction?

We determined the extent to which distributed patterns across space (ROIs) and time contribute to clip prediction by benchmarking the LSTM classifier against competing alternatives. First, we classified based on inputs at each time step, *x*_*t*_, using a feed-forward network consisting of several fully-connected layers (FF classifier). They were able to classify at no more than 51.03% accuracy (5 layers).

As FF classifiers do not capture short-term temporal relationships, we next employed temporal convolutional networks (TCN classifier). We used a kernel width (*W*) of 10 time steps, such that the number of parameters of the TCN classifier closely matched that of the LSTM (see Section 2.5 for kernel details). Classification accuracy was only 65.04%. Only with 5 times the number of parameters of the LSTM did the TCN classifier (*W* = 50) even approach LSTM performance at 83.26%. Together, the results reveal that spatiotemporal patterns are distributed across time and are most effectively captured by LSTMs capable of capturing long-term dependencies.

Finally, if capturing temporal information and long-term dependencies are important for classification, we reasoned that performance should be affected by temporal order. To test this, we shuffled the temporal order of each clip and then trained an LSTM classifier. To preserve autocorrelation structure in fMRI data while shuffling, we used a wavestrapping approach [19]. Classification accuracy reduced drastically to 64.29%.

### 3.3 Low-dimensional trajectories as spatiotemporal signatures

We sought to characterize the intrinsic dimensionality of the data required for clip prediction. High-dimensional LSTM outputs were projected to a lower dimensional space via the DR-FC layer (Fig. 1B). For visualization purposes, we projected to 3 dimensions. Despite the drastic dimensionality reduction of the LSTM outputs, classification accuracy was 77.30% (compared to 87.35% for 150 dimensions). In Fig. 3A, the low-dimensional outputs are shown for each clip, which we refer to as *trajectories* (consecutive low-dimensional 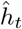 states, (*x*(*t*)*, y*(*t*)*, z*(*t*)), describe the trajectory in “state space”). Emphasizing that high-dimensional LSTM states are important to capture temporal properties, note that the size of the hidden state layer was kept at *N*_*h*_ = 150 (Fig. 1B); e.g., setting *N*_*h*_ = 3 reduced the classification performance drastically to less than 50%.

**Figure 3:**
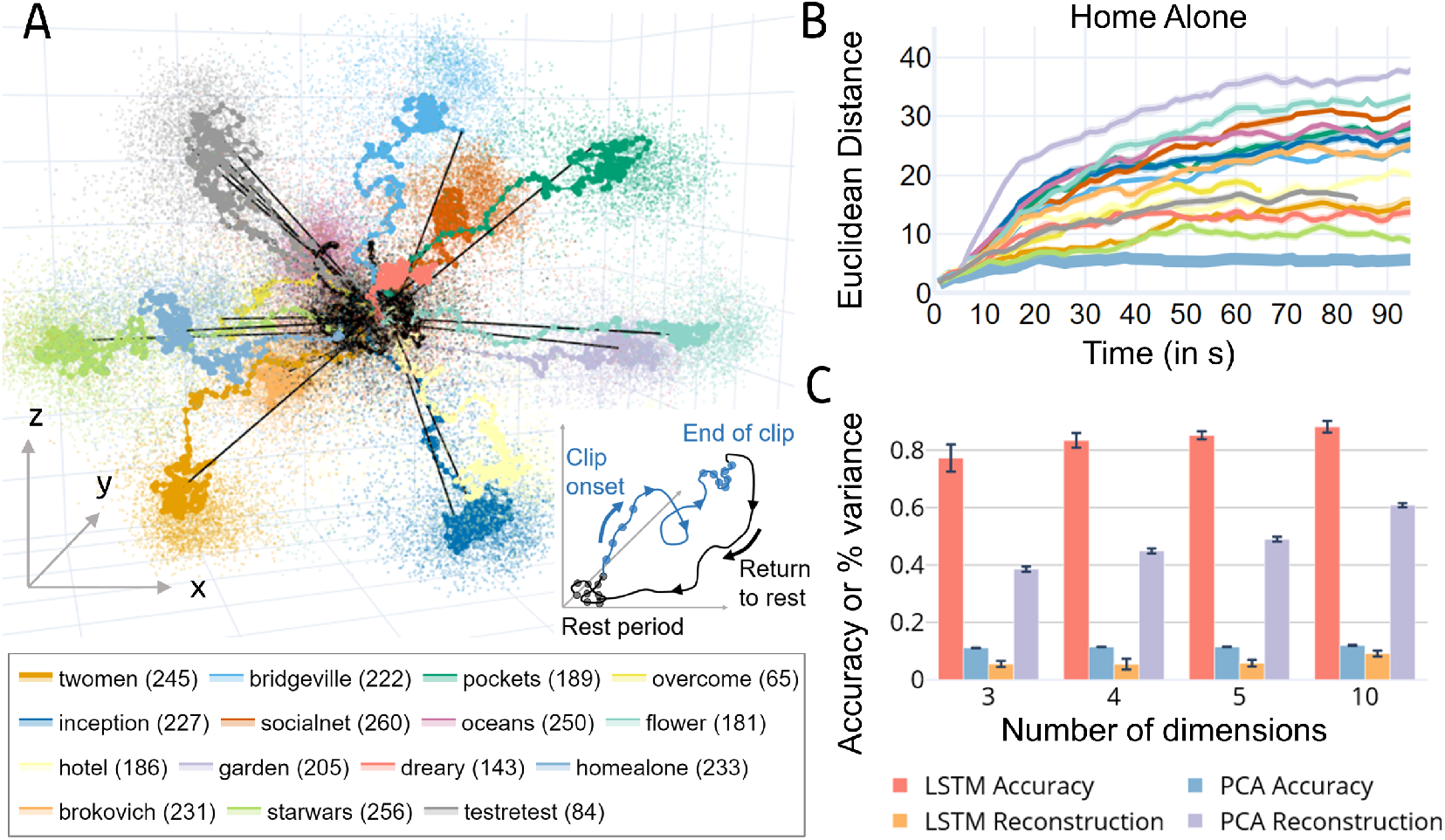
Low-dimensional trajectories. (A) Trajectories for all clips. The inset illustrates how to interpret the main result: for each clip time flows outward until the end of the clip. Solid line: mean trajectory averaged across participants; scattered points: projections for each participant at a given point in time. Trajectory associated with rest periods between clips are shown in black. (B) Distance between trajectories. The distance between the clip trajectory while watching *Home Alone* and the mean trajectory across participants for a second clip was computed. Thicker line corresponds to the distance of participants’ *Home Alone* trajectories to the mean of this clip. (C) Clip prediction accuracy and fraction of variance captured after reconstruction using low-dimensional models.

To visualize a notion of proximity between trajectories, we computed Euclidean distance between them as a function of time (Fig. 3B). To compute the distance of clip *A* from *B*, we first computed the mean trajectory of *B* averaged across participants. For each participant’s clip A trajectory, we computed Euclidean distance from the mean trajectory of *B* at every time step. Note that the proximity of a clip from itself is not zero (indicated by a thicker line), and is a measure of the consistency of participant trajectories around the clip’s mean. The evolution of trajectories closely matched the temporal accuracy obtained using the original LSTM classifier. Clip trajectories were initially close-by, but slowly separated during the first minute of the clip.

Performance with low-dimensional encoding was surprisingly high; 3 dimensions yielded 77.30% accuracy. We further investigated low-dimensional projections with 4, 5, and 10 dimensions, at which point performance (87.23%) was very similar to that of full dimensionality. These results reveal that latent representations with as few as 10 dimensions capture essential discriminative information.

However, the dimensionality-reduction fully-connected (DRFC) layer was essential in capturing this information. For reference, application of standard PCA on the input data yielded prediction accuracy at substantially lower values (Fig. 3C). Note that the latent space uncovered by the LSTM encoder, which was successful at movie prediction even with as few as 10 dimensions, captured informational content of the fMRI time series that was substantially distinct from the fMRI signal itself. To appreciate this, consider that using the LSTM decoder (Fig. 1C), to reconstruct the input time series, 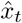, from the projections of LSTM states, 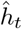, only captured a very modest amount of signal variance (less than 10%). Again, for comparison the same was done with PCA; not surprisingly, reconstructed signals recovered up to 60%.

### 3.4 Spatiotemporal saliency maps

The representations that support classification lie in an abstract space that is disconnected from the original brain activation signals. However, it is important to uncover how brain regions contribute to classification as a function of time. To do so, we used saliency maps (see Section 2.4). The majority of salient inputs occurred within the first 90 seconds of the clip, paralleling the increase in classification accuracy. After this period, changes at the input did not cause sizeable changes to the class score.

To assess the contribution of different brain networks (see legend in Fig. 4) to saliency, we averaged ROI saliency within each subnetwork to compute “network saliency”, as shown in Fig. 4A (left) for the *Inception* clip. To evaluate the evolution of saliency across time, we time-averaged ROI and network saliency for each 10-second non-overlapping window. We observed fluctuations in relative network contributions across time during the initial segment of the clip (illustrated here up to 40 s). At the level of brain regions, it is noteworthy that several ROIs with high saliency at the beginning of the clip were not captured by the time-averaged saliency map.

**Figure 4:**
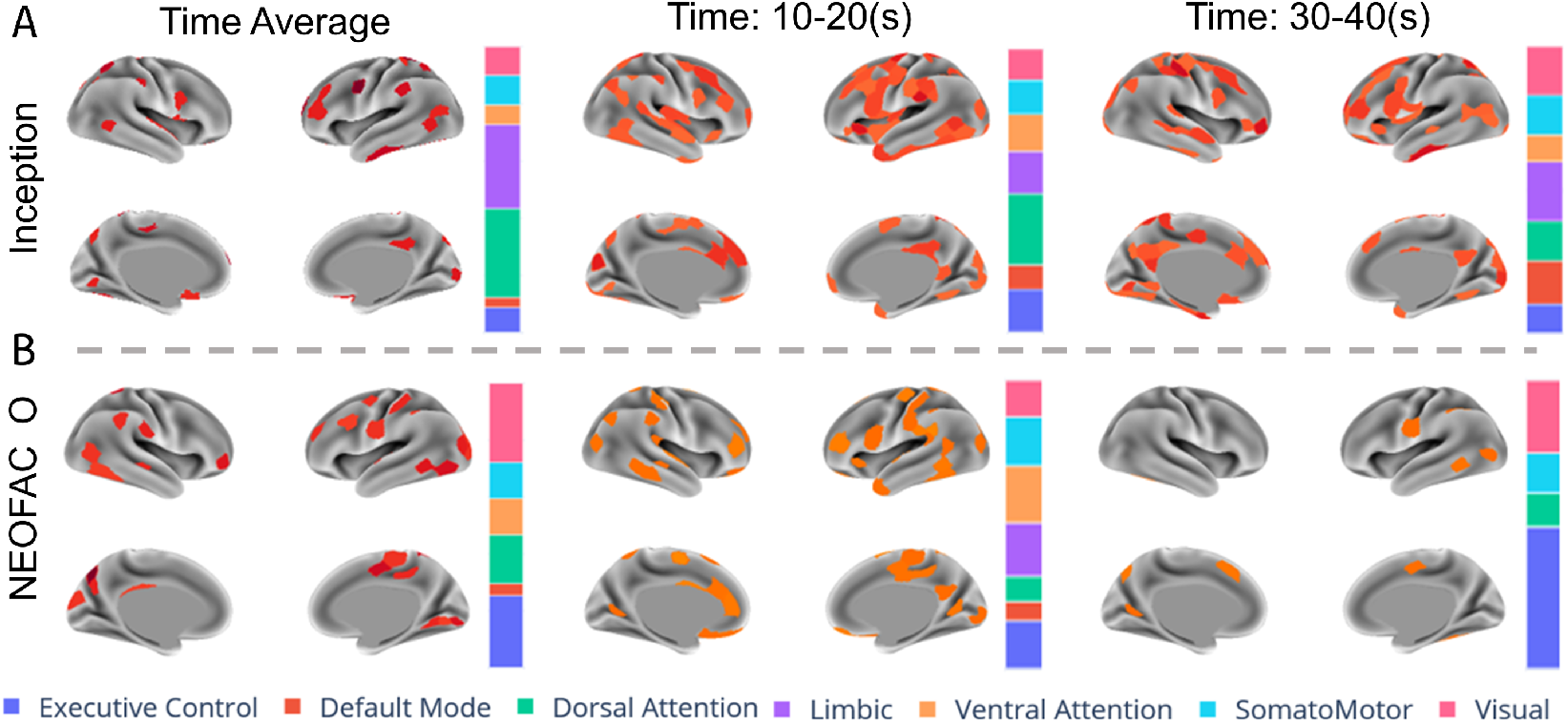
Saliency maps. (A) Top-30 most salient brain regions for predicting *Inception*. On the left, saliency was averaged across time. The right two sets of maps show mean saliency in two 10-second windows. (B) Top-30 most salient regions for predicting *openness to experience* scores (NEOFAC O). Brain maps were again based on *Inception*, the best clip for predicting the trait. The normalized contributions of 7 networks to saliency are shown alongside brain maps (colors correspond to networks and height to %).

### 3.5 Predicting behavior and personality

In recent years, researchers have attempted to employ brain data to predict a participant’s behavioral capabilities, as well as personality-based measures [20, 21, 22, 23, 24]. We hypothesized that spatiotemporal information captured by the LSTM architecture would provide valuable predictive information. The HCP dataset includes an extensive evaluation of each participant conducted outside the scanner. Here, we targeted available scores for fluid intelligence, verbal IQ, as well as scores associated with the NEO Five-Factor Inventory (with dimensions openness to experience (O), conscientiousness (C), extraversion (E), agreeableness (A), and neuroticism (N)).

Recent work has shown that participants with high scores along particular behavior/personality dimensions have similar brain responses to naturalistic stimuli; in contrast, low scorers had more idiosyncratic responses that were less similar to both high scorers and other low scorers [25]. Accordingly, here we focused on data from participants with scores at the top and bottom 10% of scores. Based on this criterion, models were trained for each behavior/personality (approx. 40 participants) separately, and generalizability was assessed using a 10-fold nested cross-validation instead of a held-out test set due to limited data. We hypothesized that participants with particular traits would generate brain states that emphasize their personality during brief clip segments, given specific stimuli during those segments (e.g., negative or positive scenes). Accordingly, we anticipated that prediction accuracy would vary across time. We thus determined optimal hyperparameters as well as the time step when prediction accuracy was highest, *t*_best_, for each clip based on the inner cross-validation. We report accuracy at *t*_best_ based on the outer cross-validation (Fig. 5).

**Figure 5:**
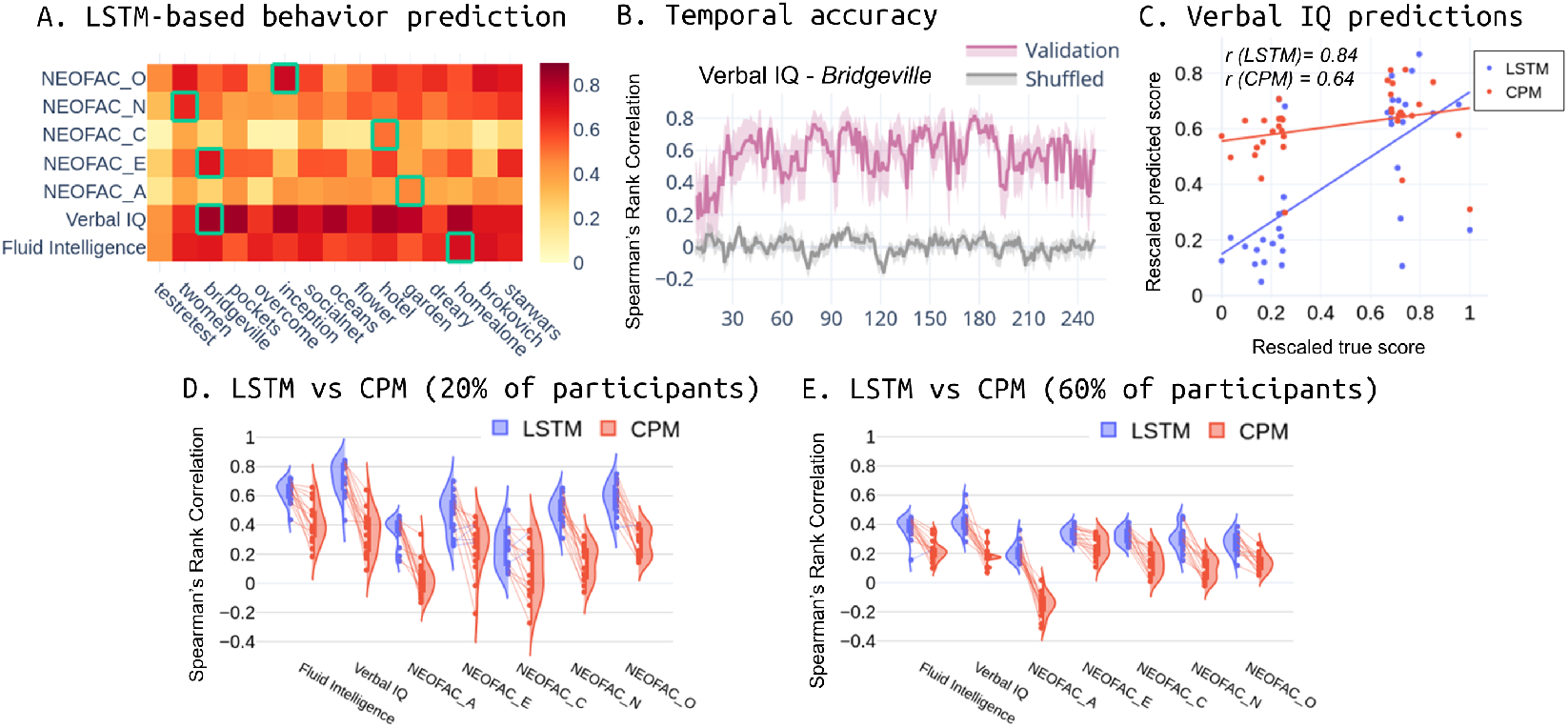
Prediction of behavior and personality. (A) Behavior prediction accuracy using LSTMs (best clip with green box). (B) Temporal accuracy for Verbal IQ while watching *Bridgeville* (best clip) along with mean null accuracy obtained by randomly shuffling behavioral measures across participants. (C) Correlation (*r*) between predicted and true verbal IQ scores (rescaled between 0 and 1) while watching *Bridgeville*. (D) Comparison of LSTM prediction accuracy with connectome-based predictive modeling (CPM) using top/bottom 10% of the scorers. Connecting lines indicate accuracy for the same clip. (E) Same as (D) but using top/bottom 30% of the scorers.

Existing techniques for predicting behavior and personality traits use what is termed *functional connectivity* as inputs to regression models [20, 22]. Functional connectivity refers to the correlation between brain regions, typically the Pearson correlation between their time series. We compared our approach to connectome-based predictive modeling (CPM; [26]), possibly the state-of-the-art in this regard. In this approach, a functional connectivity matrix is initially formed for each clip based on all ROIs. Subsequently, the entries in the matrix (or edges) that correlate with the behavioral measure beyond a set threshold (here, 0.2) are retained and a linear model is fit to predict behavioral or personality scores based on these edge weights (i.e., the functional connectivity). Unlike LSTMs, predictions are made based on the entire clip, and not at each time step.

We used all clips for training (rather than training a separate model based on each clip) to promote learning representations that are not idiosyncratic to a particular clip, and thus generalizable across clips. Nevertheless, model performance, measured as Spearman’s rank correlation between predicted and measured scores, was computed for each clip to understand how behavior/traits were captured better by certain stimuli. LSTM accuracy varied across behavior/personality measures and clips, but was consistently and robustly higher than CPM (Fig. 5A and D). Note that chance performance for each measure based on permutation testing (1000 iterations; [26]) resulted in correlations of less than 0.1. We illustrate the correlation between predicted and true scores in the case of verbal IQ (Fig. 5C). To ascertain that the improved performance of LSTM was not driven by the selection of the 10% cutoff, we increased the number of participants to include the top/bottom 30% of the scorers. Again, LSTMs performed considerably better than CPM, although prediction accuracy decreased for both models (Fig. 5E). To illustrate the temporal dimension of performance, we used the *Bridgeville* clip to predict Verbal IQ (Fig. 5B). As observed with most clips (not shown), accuracy increased at the beginning of the clip but continued to fluctuate. The results support the idea that behavior and personality traits are best captured during specific clip segments.

Finally, we sought to determine the importance of brain regions and brain networks to prediction via saliency maps (Fig. 4B). For prediction of openness to experience scores based on the *Inception* clip, the time-averaged saliency map across the entire clip (left) did not capture several salient regions at other time windows, again illustrating the temporal dimension of saliency maps. Finally, it is noteworthy that brain regions involved in classification of the *Inception* clip (Fig. 4A) were quite different from those involved in predicting the openness personality trait.

## 4 Discussion

We employed an LSTM-based architecture to characterize distributed spatiotemporal patterns of dynamic fMRI data. Representations associated with clips required capturing long-term dependencies, were consistent across participants, and generalized to previously unseen participants. Low-dimensional trajectories obtained using LSTMs captured important temporal properties in few dimensions and served as “signatures” for movie clips. Saliency maps uncovered brain regions and their time-varying importance to prediction. While recent work has utilized relatively static characterizations of fMRI data (e.g., functional connectivity) to predict individual differences in behavior and personality, here we show that spatiotemporal patterns adds substantial predictive information. Across a range of behavioral and personality measures, LSTMs outperformed the state-of-the-art. The results hint that neural responses are composed of stimuli- and individual-related components. Understanding their interaction and intrinsic dimensionality is a fruitful avenue for future research. Overall, our work provides evidence that brain dynamics must be embraced for a fuller characterization of underlying processes. Finally, whereas our results pertain to fMRI data, our LSTM framework can readily be applied to other types of brain data for discovering and interpreting brain dynamics.

### Broader Impact

Experiments in fMRI are typically tightly-controlled and analyzed using univariate approaches that explain how various regions are linked to behavior. Whereas such experiments have provided valuable insights into how the brain functions, more “naturalistic” stimuli likely engage the brain in other ways. For a fuller understanding of processes that support mental functions, we argue that brain dynamics must be addressed head-on. However, there is a relative dearth of methods to study dynamics. Here, we propose the use of recurrent neural network architectures to characterize dynamics in brain data. Our work shows that to relate neural responses to the stimuli that evoke them, it is essential to characterize distributed spatiotemporal patterns in these responses. Furthermore, individual differences in such spatiotemporal patterns were predictive of participants’ behavioral and personality characteristics. Our work provides promising results that suggest the existence of complex but consistent spatiotemporal structure in brain data obtained with fMRI. Application of these techniques to fMRI and other types of brain data obtained during naturalistic and dynamic experimental paradigms can advance our understanding of how the brain integrates diverse processes to support mental functions.

## Acknowledgments and Disclosure of Funding

M.V. was supported by a fellowship by the Brain and Behavior Initiative, University of Maryland, College Park. L.P. is supported by the National Institute of Mental Health (R01 MH071589 and R01 MH112517). The authors would like to thank Joseph Barrow and Vaishnavi Patil for helpful discussions. Data were provided [in part] by the Human Connectome Project, WU-Minn Consortium (Principal Investigators: David Van Essen and Kamil Ugurbil; 1U54MH091657) funded by the 16 NIH Institutes and Centers that support the NIH Blueprint for Neuroscience Research; and by the McDonnell Center for Systems Neuroscience at Washington University.

## Footnotes

1 Code available at https://github.com/makto-toruk/lstm-fmri-dynamicstrajectories.

